# A secondary β-hydroxybutyrate metabolic pathway linked to energy balance

**DOI:** 10.1101/2024.09.09.612087

**Authors:** Maria Dolores Moya-Garzon, Mengjie Wang, Veronica L. Li, Xuchao Lyu, Wei Wei, Alan Sheng-Hwa Tung, Steffen H. Raun, Meng Zhao, Laetitia Coassolo, Hashim Islam, Barbara Oliveira, Yuqin Dai, Jan Spaas, Antonio Delgado-Gonzalez, Kenyi Donoso, Aurora Alvarez-Buylla, Francisco Franco-Montalban, Anudari Letian, Catherine Ward, Lichao Liu, Katrin J. Svensson, Emily L. Goldberg, Christopher D. Gardner, Jonathan P. Little, Steven M. Banik, Yong Xu, Jonathan Z. Long

## Abstract

β-hydroxybutyrate (BHB) is an abundant ketone body. To date, all known pathways of BHB metabolism involve interconversion of BHB and primary energy intermediates. Here we show that CNDP2 controls a previously undescribed secondary BHB metabolic pathway via enzymatic conjugation of BHB and free amino acids. This BHB-ylation reaction produces a family of endogenous ketone metabolites, the BHB-amino acids. Genetic ablation of CNDP2 in mice eliminates tissue amino acid BHB-ylation activity and reduces BHB-amino acid levels. Administration of BHB-Phe, the most abundant BHB-amino acid, to obese mice activates neural populations in the hypothalamus and brainstem and suppresses feeding and body weight. Conversely, CNDP2-KO mice exhibit increased food intake and body weight upon ketosis stimuli. CNDP2-dependent amino acid BHB-ylation and BHB-amino acid metabolites are also conserved in humans. Therefore, the metabolic pathways of BHB extend beyond primary metabolism and include secondary ketone metabolites linked to energy balance.

## Introduction

β-hydroxybutyrate (BHB) is an abundant mammalian ketone body.^1,2^ Levels of circulating BHB rise during ketosis, a physiologic and metabolic state that is stimulated when carbohydrate availability is low and lipolysis rates are high. Once produced, BHB functions as a metabolic fuel that can be catabolized by metabolic tissues like the brain and heart for ATP production. In addition to its role as a metabolic fuel, growing evidence also suggests that BHB can exert signaling functions by diverse molecular mechanisms, including engaging G-protein coupled receptors,^3,4^ post-translational modification of proteins,^5^ and inhibition of nuclear histone deacetylases.^6^ In recent years, systemic elevation of BHB via fasting, prolonged exercise, or nutritional ketosis has attracted considerable attention for potential beneficial applications in obesity,^7^ diabetes,^8^ cancer,^9^ and other age-associated chronic diseases.

The classical enzymatic pathways for hepatic BHB production (ketogenesis) and extrahepatic BHB catabolism (ketolysis) are well-established. In the liver, fatty acid oxidation leads to the production of acetyl-CoA, which, via the sequential enzymatic action of HMGCS2, HMGCL, and BDH1, ultimately results in biosynthesis of BHB.^10^ Once produced, BHB is exported via monocarboxylate transporters (MCTs) to the circulation.^11^ BHB can then be taken up into metabolic tissues, including muscle, heart, and brain, where it is subsequently catabolized into acetyl-CoA and used for ATP production.^12^ A growing number of recent reports also indicate enzymes of ketogenesis and ketolysis operate in a number of additional cell types. For instance, BHB production has also been reported in PRDM16+ adipocytes,^13^ gut Lgr5+ intestinal stem cells,^14^ and certain T cells.^15^ Notably, to date all of these pathways involve the same biochemical interconversion of BHB to primary metabolic intermediates that are directly used for ATP production; whether BHB can be further derivatized by additional biochemical pathways has not been thoroughly explored.

Previously, Jansen *et al.* showed that CNDP2 catalyzes the condensation of lactate and amino acids in vitro,^16^ and we established this biochemical pathway to be a physiologically relevant synthetase reaction in vivo.^17^ The most abundant member of the N-lactoyl amino acids, Lac-Phe (N-lactoyl-phenylalanine), is an exercise- and metformin-inducible metabolite that suppresses food intake.^18,19^ CNDP2 is highly expressed in anatomically diverse immune cells, as well as epithelial cells of the gut and kidney, rather than classical tissues associated with lactate metabolism such as muscle or liver.^20^ These data show that the enzymatic pathways of lactate metabolism extend beyond glycolysis and primary metabolism, and include secondary metabolic pathways that produce lactate-derived signaling metabolites.

Despite many differences in their physiologic regulation and function, BHB and lactate exhibit a high degree of chemical similarity. First, BHB and lactate are structurally similar hydroxycarboxylic acids that differ by only a single methylene. Second, BHB and lactate are both transport substrates for the MCTs, demonstrating that their structural similarity also translates to a functional similarity in molecular recognition, at least with respect to the active site of certain solute carriers. Lastly, both BHB and lactate can reach high, millimolar levels in the blood following specific physiologic conditions (sprint exercise for lactate and ketosis for BHB). Based on these observations, we considered the possibility that CNDP2, like the MCTs, might also exhibit substrate promiscuity and accept BHB as a substrate. This postulated CNDP2-dependent BHB-ylation of free amino acids would represent a previously unknown pathway of BHB metabolism and produce a class of orphan metabolites, the BHB-amino acids (N-β-hydroxybutyryl amino acids). However, we were unable to find any evidence for enzymatic BHB-ylation of free amino acids in the published literature. We also were unable to find prior annotation of any BHB-amino acids as endogenous metabolites in public databases such as METLIN,^21^ HMDB,^22^ or GNPS.^23^

Here we show that CNDP2-dependent BHB-ylation of free amino acids is indeed a physiologic and previously unrecognized enzymatic reaction in vitro and in vivo. The product of this biochemical reaction, the BHB-amino acids, are endogenously present in mouse and human plasma and exhibit ketosis-inducibility and genetic regulation. BHB-Phe is the most abundant BHB-amino acid and a structural and functional congener of Lac-Phe. Conversely, CNDP2-KO mice exhibit increased food intake and body weight on ketogenic diet or following ketone ester administration. Lastly, CNDP2-mediated amino acid BHB-ylation and the BHB-amino acid metabolites are conserved in humans. These data establish a secondary ketone metabolic pathway linked to energy balance.

## Results

### CNDP2 catalyzes BHB-ylation of amino acids in vitro

Based on the chemical similarity between lactate and BHB, we hypothesized that CNDP2 might exhibit enzyme active site promiscuity and, in addition to lactate^17^, also accept BHB as a condensation substrate (**Fig. 1A**). To experimentally determine whether CNDP2 can catalyze the BHB-ylation of phenylalanine in vitro, we incubated BHB and phenylalanine (20 mM each) with cell lysates from HEK293T cells that were transiently transfected with flag-tagged mouse CNDP2 or GFP control. Overexpression of CNDP2 protein was confirmed by Western blotting (**Fig. S1A**). After incubation at 37°C for 1 h, we used liquid chromatography-mass spectrometry (LC-MS) to measure the expected condensation product BHB-Phe (N-β-hydroxybutyryl phenylalanine). As shown in **Fig. 1B**, CNDP2-transfected cell lysates exhibited >140-fold greater phenylalanine BHBylation activity compared to GFP-transfected cell lysates. CNDP2 can therefore synthesize BHB-Phe, in addition to its previously reported Lac-Phe synthesis activity. We performed several control experiments to examine the substrate specificity of the CNDP2-dependent BHB-ylation reaction. CNDP2 exhibited similar rates of amino acid BHB-ylation and N-lactoylation, but could not accept shorter (e.g., acetate, C2) or longer (e.g., octanoate, C8) organic acids as substrates (**Fig. 1C**). On the amino acid side, CNDP2 exhibited the fastest BHB-ylation activity using phenylalanine as a substrate (**Fig. 1D**). Some other hydrophobic amino acids were also accepted but with lower activity (<20%) compared to phenylalanine, and no activity was observed with many of the amino acids tested (**Fig. 1D**). CNDP2 did not exhibit BHB-ylation of the free N-terminus of a peptide substrate (**Fig. S1B**). CNDP2 was also unable to accept the BHB isomer 3-hydroxisobutyric acid (3-HIB) as a substrate, though we did observe minor CNDP2 condensation activity with 2-hydroxybutyrate (2HB) (**Fig. S1C**).^24,25^

**Fig. 1.**
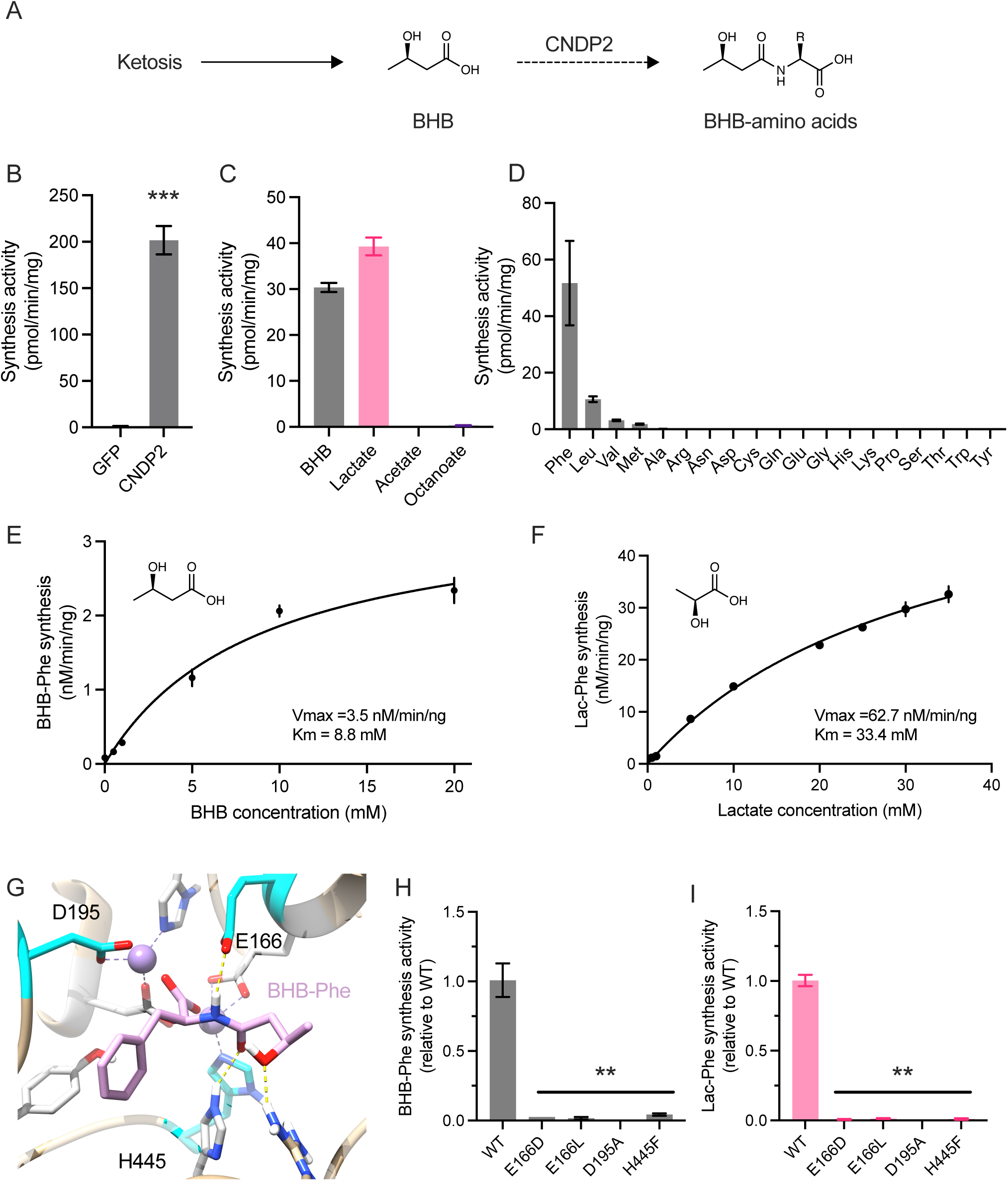
CNDP2 catalyzes amino acid BHB-ylation in vitro. (A) Schematic of CNDP2-dependent Lac-Phe synthesis and the proposed CNDP2-dependent BHB-Phe synthesis reaction. (B) BHB-Phe synthesis activity of cell lysates transfected with GFP or mouse CNDP2-flag (C,D) Synthesis activity of cell lysates transfected with GFP or mouse CNDP2-flag and incubated with the indicated monocarboxylate with Phe (C) or the indicated amino acid with BHB (D) (E,F) Michaelis-Menten kinetics of recombinant purified mouse CNDP2-flag protein with BHB (E) or lactate (F) as the organic acid donor. (G) Molecular docking of BHB-Phe into the mouse CNDP2 active site. (H,I) BHB-Phe synthesis activity (H) or Lac-Phe synthesis activity (I) of HEK293T cells transfected with the indicated mouse WT or mutant CNDP2 plasmid. For enzyme assays, organic acids and amino acids were incubated at a concentration of 20 mM at 37°C for 1 hour. For B-I, N=3-4/group. Data are shown as means ± SEM. P-values were calculated by Student’s two-sided t-test.

To quantitatively compare the kinetics of CNDP2-dependent BHB-ylation and N-lactoylation, we generated purified recombinant mouse CNDP2-flag protein for in vitro kinetic assays. The resulting kinetic data, which were fit to Michaelis-Menten kinetics, revealed that BHB is in fact a higher affinity substrate for the CNDP2 active site than lactate (Km for BHB = 8.8 mM; Km for lactate = 33.4 mM, **Fig. 1E,F**). By contrast, CNDP2-dependent Lac-Phe synthesis is faster than that of BHB-Phe synthesis under conditions of saturating substrate concentrations (Vmax for lactate = 62.7 nM/min/mg, Vmax for BHB = 3.5 nM/min/mg, **Fig. 1E,F**).

Lastly, we docked the product BHB-Phe into the CNDP2 active site (**Fig. 1G** and see **STAR Methods**). This modeling predicted several interactions of BHB-Phe with key active site residues, including E166, D195, and H455. We generated single point mutations of each of these residues by transient transfection to HEK293T cells (**Fig. S1D**). In each case, mutations of any of these active site residues concomitantly reduced both CNDP2-dependent BHB-Phe and Lac-Phe production (**Fig. 1H,I**). Therefore BHB-ylation and N-lactoylation activities are both entirely encoded within the CNDP2 polypeptide. In addition, these two catalytic activities cannot be readily dissociated by single point mutations in the active site pocket.

### CNDP2-dependent BHB-ylation in mouse tissues

To determine if endogenous CNDP2 catalyzes amino acid BHB-ylation in mouse tissues, we first examined the tissue expression of CNDP2 by Western blot using an anti-CNDP2 antibody. The specificity of this antibody was confirmed using tissues from CNDP2-KO mice (**Fig. S2A**). CNDP2 protein levels were highest in kidney and gut, and lower in many of the other tissues examined (**Fig. 2A**). Next, we performed in vitro BHB-Phe synthesis activity using crude total lysates of kidney, gut, brain, liver, or quadriceps tissues from WT mice (see **STAR Methods**). We selected these tissues because of their wide range of CNDP2 protein expression. As shown in **Fig. 2B**, the highest BHB-Phe synthesis activity was observed in kidney and gut (∼10-30 pmol/min/mg), while brain, liver, and quadriceps exhibited lower, but detectable BHB-Phe synthesis activity (∼1-5 pmol/min/mg). This pattern of BHB-Phe synthesis across tissues largely paralleled the protein expression of CNDP2 in these same tissues. In addition, the temperature dependence of the renal BHB-Phe synthesis activity also paralleled that of recombinant CNDP2 protein (**Fig. S2B,C**). Therefore the tissue expression pattern and temperature profile of CNDP2 protein both correlate with the tissue BHB-Phe synthesis activity.

**Fig. 2.**
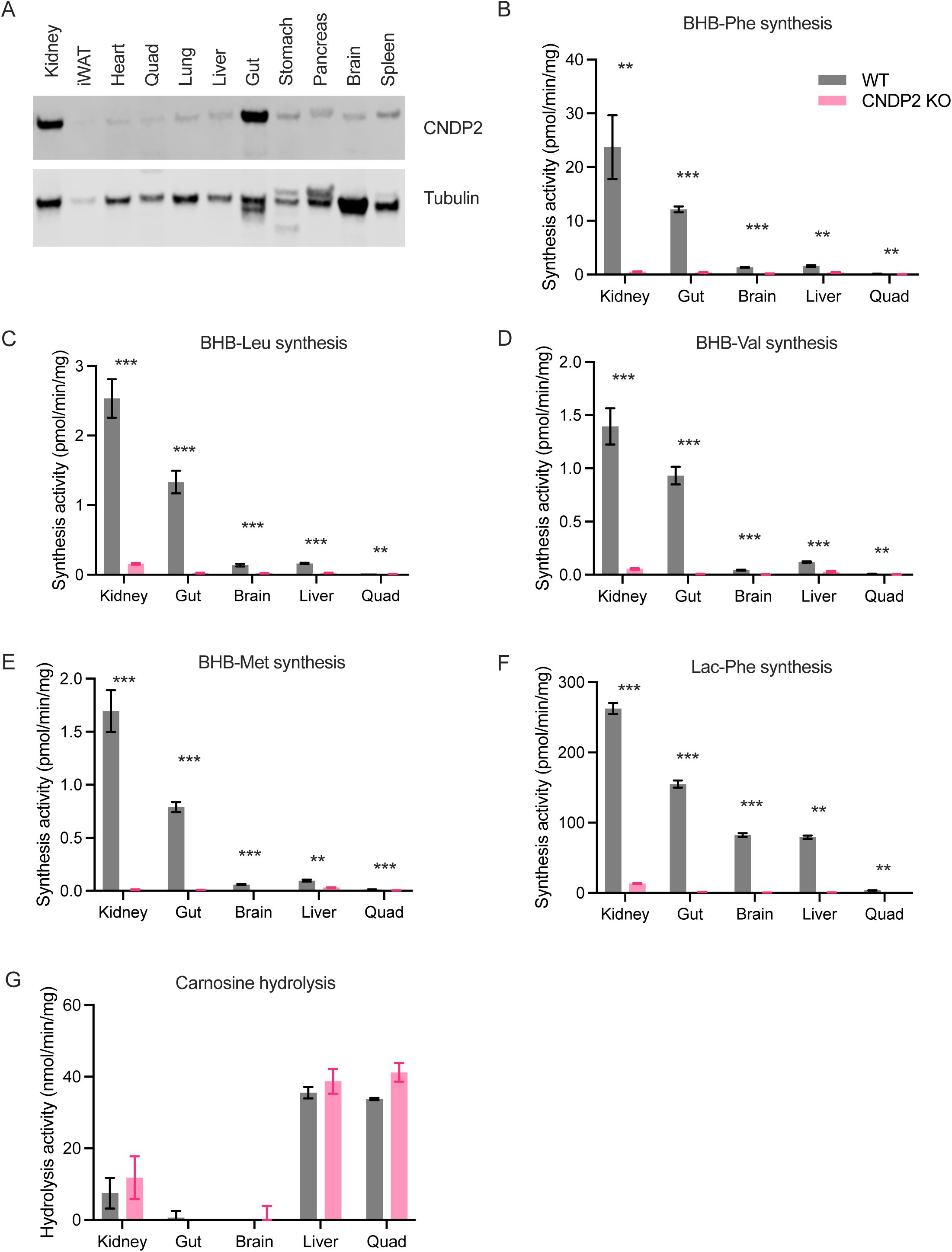
CNDP2 is the principal BHB-amino acid synthetase in mouse tissues. (A) Western blot of the indicated mouse tissues using an anti-CNDP2 (top) or anti-tubulin (bottom) antibody. (B-G) Enzyme activities of tissues from WT or CNDP2-KO mice when provided with BHB and Phe (B), BHB and Leu (C), BHB and Val (D), BHB and Met (E), lactate and Phe (F), or carnosine (G) as substrates. For enzyme assays, organic acids and amino acids were incubated at a concentration of 20 mM at 37°C for 1 hour. For B-G, N=3-4/group. Data are shown as means ± SEM. P-values were calculated by Student’s two-sided *t*-test.

We used tissues from CNDP2-KO mice to determine the contribution of CNDP2 to the tissue BHB-Phe synthesis activity. As shown in **Fig. 2B**, both kidney and gut BHB-Phe synthesis activity was largely abolished (>95%) in tissues from CNDP2-KO mice. The smaller BHB-Phe synthesis activity in other tissues was also greatly diminished (>85% reduced in brain, >75% reduced in liver, and >60% reduced in quadriceps). Using leucine, valine, methionine as substrates for in vitro BHB-ylation, a similar pattern of BHB-Leu, BHB-Val, and BHB-Met synthesis across WT and CNDP2-KO tissues was observed (**Fig. 2C-E**): kidney and gut tissues from WT mice both exhibited the highest BHB-amino acid synthesis activity, and this activity was largely abolished in tissues from CNDP2-KO mice. In additional control experiments, we assayed tissue Lac-Phe synthesis activity, which again exhibited a similar pattern and CNDP2-dependence to that of BHB-amino acid synthesis (**Fig. 2F**). By contrast, carnosine hydrolysis across tissues exhibited a distinct pattern with highest activity in liver and quadriceps and little activity in the kidney, gut, and brain (**Fig. 2G**). Importantly, the carnosinase activity was not altered in CNDP2-KO tissues (**Fig. 2G**). We conclude that CNDP2 is the principal enzyme responsible for BHB-amino acid synthesis activity in mouse tissues. In addition, despite its previously annotated in vitro activity, CNDP2 is not a major tissue carnosinase.

### BHB-amino acids are endogenous mouse metabolites

The product of the CNDP2-catalyzed BHB-ylation reaction, BHB-amino acids, have not been previously reported as endogenous metabolites. We therefore developed a targeted metabolomics approach to determine if BHB-amino acids can be detected in mouse plasma. We first synthesized an authentic BHB-Phe standard by classical amide coupling between BHB and phenylalanine (**Fig. 3A** and see **STAR Methods**). Fragmentation of the authentic BHB-Phe standard revealed a major daughter ion corresponding to Phe (m/z = 164) and a second, smaller daughter ion corresponding to the decarboxylation product (m/z = 206). Next, we developed a targeted multiple reaction monitoring (MRM) method on a high-performance liquid chromatography coupled to triple quadrupole mass spectrometry (QQQ-LC/MS) to monitor the parent to phenylalanine transition for BHB-Phe. In mouse plasma, we identified an endogenous peak that eluted at an identical retention time with the authentic standard (**Fig. 3A**). To exclude the possibility that this method may also be detecting isobaric 2-hydroxybutyrate (2-HB)-and 3-hydroxyisobutyrate (3-HIB)-phenylalanine isomers, we synthesized authentic standards of 2-HB-Phe and 3-HB-Phe and developed MRM methods that could distinguish between each of the three molecules. While 2-HB-Phe and 3-HIB-Phe also yielded daughter ions corresponding to phenylalanine, we also identified unique transitions for each of these isomers (2-HB-Phe: 260>102, fragmentation at N-Cα; 3-HIB-Phe: 250>220, loss of CH_3_O, **Fig. S3A,B**). For BHB-Phe, >99% of the total signal was detected using the 250>164 transition, 2-HB-Phe was detected using both 260>102 and 250>164 (in a 6:1 ratio), and 3-HIB-Phe was detected using both 250>220 and 250>164 (in a 1.6:1 ratio) (**Fig. S3C**). The signal from the endogenous peak was found to be comprised >99% of the 260>164 transition (**Fig. S3C**). Therefore the endogenous signal is BHB-Phe; in addition, 2-HB-Phe and 3-HIB-Phe are not endogenous metabolites.

**Fig. 3.**
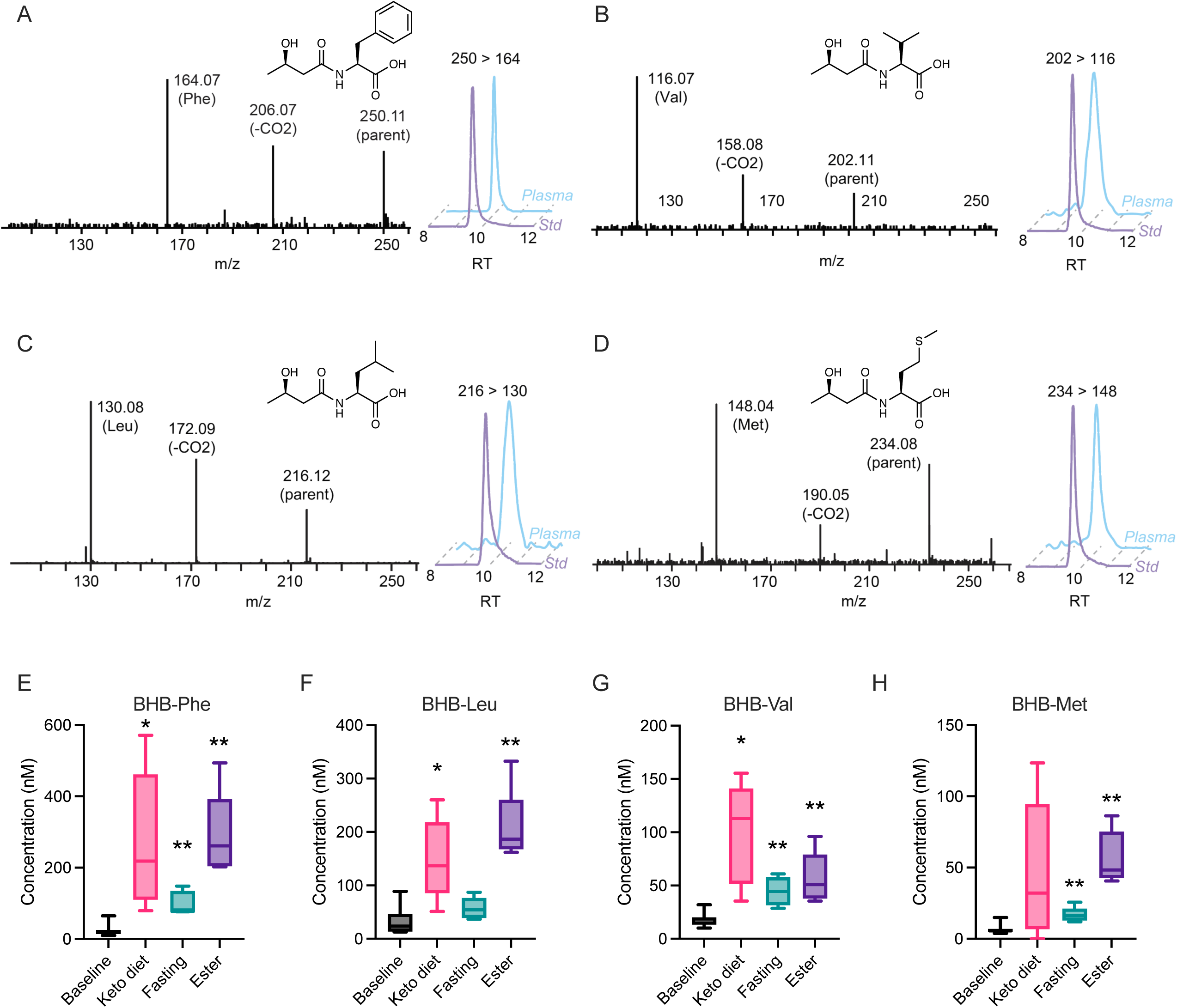
Detection and ketosis-inducibility of BHB-amino acids in mouse plasma. (A-D) Tandem mass spectrometry fragmentation of the authentic standard (left) and co-elution of the standard and the endogenous peak from mouse plasma (right) using the indicated multiple reaction monitoring transition for BHB-Phe (A), BHB-Val (B), BHB-Leu (C), and BHB-Met (D). (E-H) BHB-amino acid quantitation in 8-9 week old male C57BL/6J mouse plasma at baseline (dark blue), after 1 week on ketogenic diet (Research Diets D21021803), after a 24 h fast (grey) or 30 min post ketone monoester drink administration by oral gavage (3 mg KE/g of body weight, light blue). For E-H, N=5/group, with the baseline N=15 (pooled from each of the three groups). Data for E-H are shown as box-and-whisker plots. P-values were calculated by Student’s two-sided *t*-test.

We next synthesized authentic standards for BHB-Leu, BHB-Val, and BHB-Met, which also exhibited the same characteristic amino acid daughter ion (**Fig. 3B-D**). Using a similar MRM approach, we also detected endogenous peaks with transition and retention time identical to that of the authentic standards (**Fig. 3B-D**), demonstrating that these other BHB-amino acids are also endogenous metabolites.

Because of their biosynthetic origin from BHB, circulating BHB-amino acids would be predicted to rise with increasing BHB levels, such as those achieved by nutritional or physiologic ketosis. Levels of BHB-amino acids in mouse blood plasma were therefore measured after one week of ketogenic diet, a 24 h fast, or oral administration of a ketone ester drink (3 g/kg of body weight). We confirmed that plasma BHB levels were elevated by each of these conditions (Fig. **S3D**). All these stimuli consistently produced robust 2-10-fold elevations in each of the BHB-amino acids (**Fig. 3E-H**). Ketogenic diet produced greater variation in induction of BHB-amino acid levels, which may reflect the more chronic nature of this perturbation (1 week) compared to the two other acute ketosis stimuli (≤ 24 h). The levels of phenylalanine, Lac-Phe and lactate were not consistently changed across the three ketosis stimuli (**Fig. S3D**). Tissue levels of BHB-Phe were also detectable and elevated after ketone ester oral gavage (**Fig. S3E**). We conclude that BHB-amino acids are endogenous, ketosis-inducible mouse metabolites.

### Genetic regulation of BHB-amino acids

To understand the genetic and enzymatic pathways that regulate circulating BHB-amino acids in vivo, we used two genetically modified mouse models in which enzymes involved in the BHB-amino acid biosynthesis or ketogenesis were genetically ablated (**Fig. 4A**). First, to test the in vivo role of CNDP2 as the key biosynthetic enzyme of BHB-amino acids, we measured BHB-amino acids in blood plasma from CNDP2-KO mice after a ketone ester drink challenge or after one week of ketogenic diet. After acute administration of ketone ester drink, CNDP2-KO mice exhibited >90% depletion of multiple plasma BHB-amino acids (**Fig. 4B**). Reductions in multiple BHB-amino acids was also observed in CNDP2-KO mice after one week on ketogenic diet (**Fig. 4C**). In both experiments, levels of plasma BHB, lactate, or phenylalanine were not consistently changed between WT and CNDP2-KO mice, and levels of Lac-Phe were, as expected, reduced in CNDP2-KO mice (**Fig. S4A,B**).

**Fig. 4.**
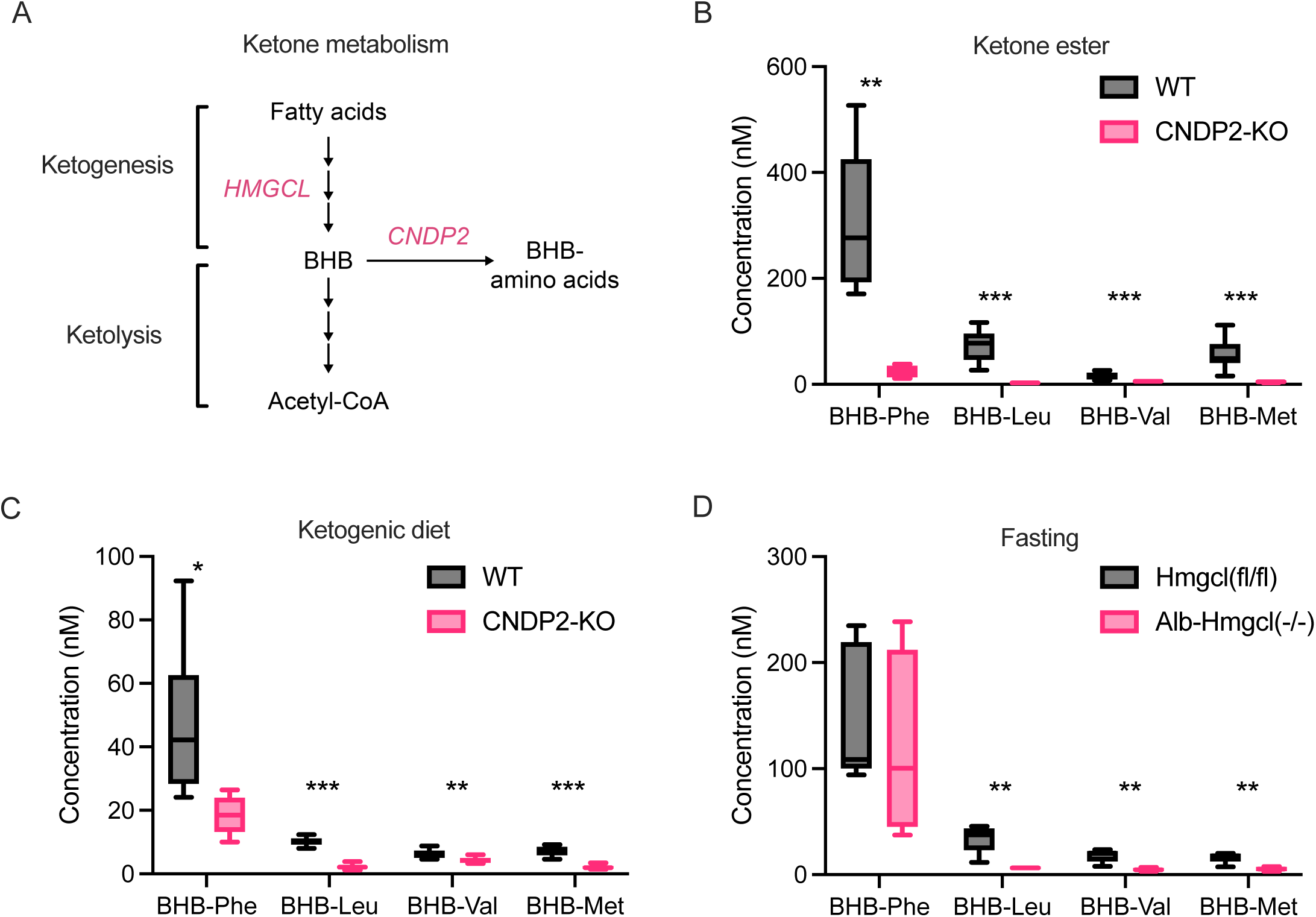
Genetic regulation of BHB-amino acids by CNDP2 and HMGCL. (A) Schematic of ketone biochemical pathways and the genetic mouse models used. (B-D) BHB-amino acid quantitation in plasma from 4-10-week-old male WT (blue) and CNDP2-KO mice (red) at 60 min post ketone monoester drink administration by oral gavage (3 mg KE/g of body weight) (B), from 7-16-week-old female WT and CNDP2-KO mice after 1 week on ketogenic diet (Research Diets D06040601, panel C), or from Hmgcl (fl/fl) vs Alb-Hmgcl(-/-) mice after a 24 h fast. For B, N = 7 for WT and 4 for KO. For C, N = 8 per group. For D, N = 5 for Hmgcl (fl/fl), N = 4 for Alb-Hmgcl(-/-). Data are shown as box-and-whisker plots. P-values were calculated by Student’s two-sided t-test.

We performed additional metabolomic profiling of organic acid-amino acid conjugates in CNDP2-KO mice. N-acetyl-Phe, N-acetyl-Val, N-acetyl-Leu, and N-acetyl-Met, as well as their corresponding free amino acids, were not changed between WT and CNDP2-KO mice (**Fig. S4C**). We were unable to detect N-propyl, N-butyryl, or N-octanoyl-amino acids (C3, C4, and C8, respectively, **Fig. S4C**), likely reflecting the low circulating abundance of the corresponding organic acids compared to acetate, lactate, and BHB.

Next, we examined the effects of liver-specific deletion of HMGCL, a critical upstream enzyme in hepatic ketogenesis (**Fig. 4A**). We obtained plasma from liver-specific knockouts of HMGCL (*Alb*-*Hmgcl*(-/-) mice) which were previously generated by crossing Albumin-cre driver mice with *Hmgcl* floxed mice.^10^ Several BHB-amino acids, such as BHB-Met, BHB-Leu, and BHB-Val, but not BHB-Phe were reduced by ∼50-80% in plasma from these animals (**Fig. 4D**). We confirmed ∼30% reductions in BHB levels in plasma from Alb-*Hmgcl*(-/-) mice and in addition found no changes in lactate, Lac-Phe, or phenylalanine levels compared to *Hmgcl* (fl/fl) controls (**Fig. S4D**). Together, these data establish the genetic and biochemical requirement for two upstream enzymes, HMGCL and CNDP2, in the regulation of circulating BHB-amino acids levels.

### BHB-Phe is a congener of Lac-Phe

BHB-Phe is the most abundant BHB-amino acid (**Fig. 3E**). This metabolite is a congener of Lac-Phe that shares both chemical similarity as well as a common biosynthetic pathway via CNDP2. We therefore considered the possibility that BHB-Phe might also be functionally similar to Lac-Phe and regulate food intake and body weight. We first used gain-of-function approaches to determine if BHB-Phe is sufficient to reduce food intake and body weight. In an initial study of DIO mice in metabolic chambers, BHB-Phe (50 mg/kg, IP) reduced food intake without affecting movement, oxygen consumption or CO_2_ production (**Fig. 5A-E**). A reduction in respiratory exchange ratio (RER) was also observed, consistent with a suppression of food intake (**Fig. 5E**). Under these conditions, plasma BHB-Phe levels peaked at ∼20 µM at the 1 hour time point and returned to baseline values by 3 h (**Fig. S5A**). In an independent experiment in home cages, we found that acute administration of BHB-Phe (50 mg/kg, IP) to DIO mice suppressed food intake without affecting water intake, establishing specific suppression of food versus fluid ingestion (**Fig. S5B**). Lastly, plasma levels of other feeding-regulating hormones, such as ghrelin, leptin, and GDF15, were also unaltered in mice following a single administration with BHB-Phe (50 mg/kg, IP, **Fig. S5C**).

**Fig. 5.**
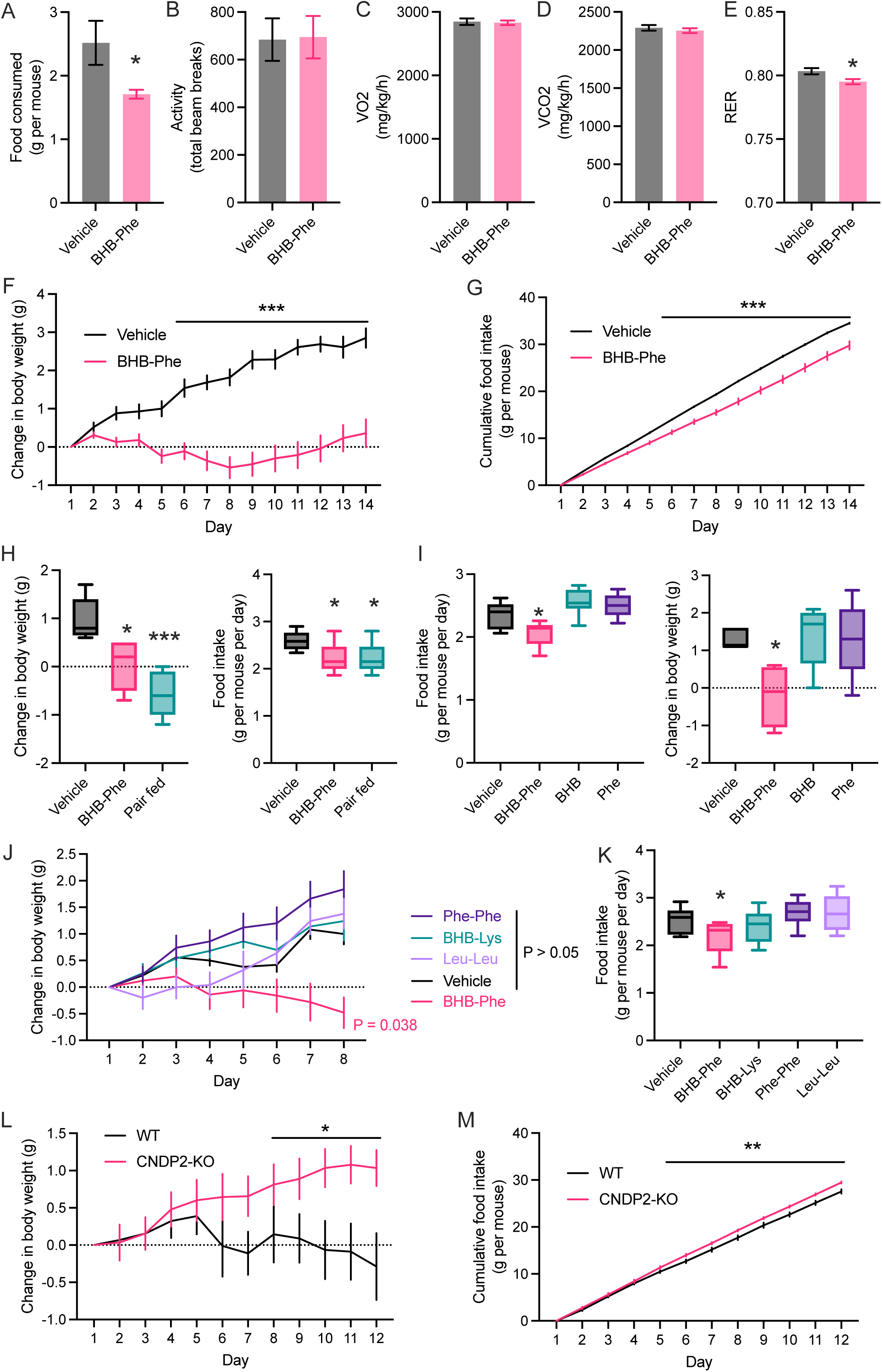
BHB-Phe suppresses food intake and body weight. (A-E) Food intake (A), ambulatory movement (B), oxygen consumption (VO_2_) (C), carbon dioxide production (VCO_2_) (D), and respiratory exchange ratio (RER) (E) of singly housed 29-week-old male DIO mice following a single injection of vehicle or BHB-Phe (50 mg/kg, IP) over a 10 h period in metabolic chambers. (F,G) Change in body weight (F) and cumulative food intake (G) of singly housed 28-week-old male DIO mice treated with vehicle or BHB-Phe (50 mg/kg/day, IP). Starting body weights were vehicle: 46.4 ± 1.4 g, BHB-Phe 45.7 ± 1.0 g (mean ± SEM). (H) Change in body weight (left) and daily food intake (right) of group housed 15-week-old male DIO mice after 6 days of treatment with vehicle, BHB-Phe (50 mg/kg/day, IP) or vehicle-treated pair-fed mice. Starting body weights were vehicle: 39.4 ± 2.3 g, BHB-Phe: 38.5 ± 0.8 g, and pair-fed: 37.8 ± 0.6 g (mean ± SEM). (I) Change in body weight (left) and food intake (right) of group housed 13-week-old male DIO mice after 9 days of treatment with vehicle, BHB-Phe, BHB, or phenylalanine (50 mg/kg/day, IP). Starting body weights were vehicle: 36.2 ± 0.7 g, BHB-Phe: 37.9 ± 2.1 g, BHB: 36.8 ± 1.1 g, Phe: 34.5 ± 0.5 g (mean ± SEM) (J,K) Change in body weight (J) and food intake (K) of group housed 14-16-week-old male DIO mice after 9 days of treatment with Phe-Phe, BHB-Lys, Leu-Leu, BHB-Phe (50 mg/kg/day, IP) or vehicle. Starting body weights were vehicle: 41.2 ± 1.3 g, BHB-Phe: 39.1 ± 1.7 g, BHB-Lys: 40.1 ± 2.2 g, Phe-Phe: 40.8 ± 1.4 g, Leu-Leu: 41.2 ± 1.3 g (mean ± SEM). (L,M) Change in body weight (L) and food intake (M) of singly housed 20-week-old male WT and CNDP2-KO mice receiving ketone esters (3 g/kg/day, PO). Starting body weights were WT: 46.8 ± 1.4 g, CNDP2-KO: 47.1 ± 1.3, P > 0.05 (mean ± SEM). For A-E, N = 7 for vehicle, N = 8 for BHB-Phe. For F,G, N = 10 per group. For H-K, N = 5 per group. For L,M, N = 9 per group. Data in A-G, J, L, and M are shown as the mean ± SEM. Data in H, I, and K are shown as box-and-whisker plots. P-values were calculated by Student’s two-sided t-test or by two-way ANOVA.

We performed chronic studies of BHB-Phe in DIO mice. Daily administration of BHB-Phe (50 mg/kg/day, IP) resulted in a durable suppression of daily food intake and, as expected, a concomitant reduction in body weight gain (**Fig. 5F,G**). At the end of the experiment, BHB-Phe-treated mice exhibited reductions in AST, ALT, and total triglycerides (TG); no changes were found in HDL- or LDL-cholesterol (**Fig. S5D**). BHB-Phe-treated mice lost the same amount of weight as pair-fed controls (**Fig. 5H**), demonstrating that the observed suppression of food intake explains the observed change in body weight in BHB-Phe-treated mice.

We performed additional control experiments to understand the structural requirements of BHB-Phe that were important for its body weight-lowering effects. First, while BHB-Phe (50 mg/kg/day, IP) efficiency suppressed body weight and food intake in DIO mice, either BHB alone or phenylalanine alone at the same doses were without effect (**Fig. 5I**). Similarly, the related metabolites BHB-Lys (50 mg/kg, IP), as well as the dipeptides Phe-Phe (50 mg/kg, IP) and Leu-Leu (50 mg/kg, IP), also failed to reduce food intake or body weight (**Fig. 5J,K**). Lastly, we tested other CNDP2-regulated BHB conjugates, including BHB-Met, BHB-Leu, and BHB-Val. These metabolites consist of BHB conjugated to hydrophobic amino acids; notably, two of them, BHB-Leu and BHB-Val, represent BHB conjugated to branched-chain amino acids (BHB-BCAAs). In this case, we observed body weight-lowering activity for all BHB conjugates tested (**Fig. S5E**). Therefore BHB-Phe and other CNDP2-regulated BHB-hydrophobic amino acid conjugates, including BHB-BCAAs, have anorexigenic and anti-obesity effects in mice, while BHB conjugates to other essential amino acids, as well as other dipeptides, do not have exhibit the same bioactivity.

Lastly, we used CNDP2-KO mice to examine the physiologic contributions of BHB-amino acids to energy balance. We previously reported a gene-by-environment interaction of the *Cndp2* gene and glycolytic stimuli: CNDP2-KO mice have normal body weights after a standard high fat diet-feeding protocol, however, upon glycolytic stimulus challenge to increase Lac-Phe levels (by treadmill exercise or by metformin treatment), knockout animals exhibit an increased food intake and body weight phenotype compared to WT controls.^17,18^ To determine the contribution of BHB-amino acids, in an initial experiment we administered ketone esters (3 g/kg/day, PO) to WT and CNDP2-KO mice that had been rendered obese by high fat diet feeding for 16 weeks. At the beginning of the experiment, body weights were not different between genotypes. By the end of the 12-day ketone ester treatment, WT mice on average lost −0.3 ± 0.4 g, while CNDP2-KO mice gained +1.0 ± 0.2 g (mean ± SEM, P < 0.05, **Fig. 5L**). CNDP2-KO mice also exhibited greater cumulative food intake than WT mice (**Fig. 5M**). In a second independent experiment, we placed WT and CNDP2-KO mice on a ketogenic diet. Initial body weights were once again not different between genotypes. By the end of the experiment, CNDP2-KO mice on ketogenic diet gained more weight and ate more food than WT mice (**Fig. S5F**). We confirmed that Lac-Phe and BHB-Phe were independently induced following sprint treadmill exercise and ketone ester or ketogenic diet treatment, respectively (**Fig. S5G**). Therefore ketone esters and ketogenic diet represent environmental perturbations that uncover the effects of *Cndp2* genotype on body weight and food intake.

### Neurobiological mechanisms of BHB-Phe

To better understand the neurobiological mechanisms by which BHB-Phe suppresses feeding, we first used pharmacological and genetic approaches to determine whether the effect of BHB-Phe might be mediated by hypothalamic melanocortin signaling, GLP1-R, or brainstem GFRAL pathways. The effect of BHB-Phe (50 mg/kg, IP) on food intake and body weight was similar in WT or MC4R-KO mice (**Fig. S6A,B**). Similarly, the GLP1-R antagonist Exendin-3 (0.1 mg/kg/day, IP) efficiently blocked the anorexigenic and anti-obesity effects of GLP-1 (2 mg/kg/day, IP, **Fig. S6C,D**); however, under these same Exendin-3 did not alter the effect of BHB-Phe on food intake and body weight (**Fig. S6E,F**). Lastly, we obtained a neutralizing anti-GFRAL antibody^26,27^ (Eli Lilly & Co. clone 8A2) and verified this antibody (10 mg/kg, SQ) completely blocked the activity of recombinant GDF15 to suppress food intake and body weight (4 nmol/kg, SQ) (**Fig. S6G**); however, the effect of BHB-Phe on feeding and body weight was unaffected by anti-GFRAL antibody administration (**Fig. S6H,I**). We conclude that the anorexigenic activity of BHB-Phe is independent of these known pathways of feeding control.

Next, we used an activity-dependent genetic labeling strategy^28^ (TRAP, targeted recombination in active populations) to identify neurons activated following pharmacological dosing of BHB-Phe. In this approach, upon BHB-Phe treatment, c-Fos-dependent recombination of a reporter cassette (tdTomato) enables permanent genetic labeling of BHB-Phe. We treated TRAP2 mice (TRAP2/Rosa26-LSL-tdTomato) with BHB-Phe (50 mg/kg, IP) followed by 4-hydroxytamoxifen 30 min later to initiate Cre-dependent recombination in BHB-Phe-activated neurons. Two weeks later, the same mice received Lac-Phe (50 mg/kg, IP) and were sacrificed 90 min later for c-Fos immunostaining (**Fig. 6A**). This experimental approach therefore enables concurrent identification of both BHB-Phe-activated (e.g., TRAP+, marked by tdTomato) and Lac-Phe-activated (e.g., c-Fos+) neurons in the same animal. We examined multiple hypothalamic and brainstem regions for TRAP+ or c-Fos+ neurons. Compared to vehicle treatment, BHB-Phe and Lac-Phe both activated neurons in multiple brain regions, including the paraventricular hypothalamic nucleus (PVH), the suprachiasmatic nucleus (SCN), the dorsomedial hypothalamic nucleus (DMH), the ventromedial hypothalamic nucleus (VMH), the arcuate nucleus of the hypothalamus (ARH), the lateral hypothalamus (LH), the lateral parabrachial nucleus (LPBN), and the nucleus of the solitary tract (NTS) (**Fig. 6B**). However, detailed analyses of TRAP+ neurons and c-Fos+ neurons in each of these regions revealed that only a small fraction (∼2-30%) of neurons were activated by both BHB-Phe and Lac-Phe, while the vast majority of activated neurons were distinct (**Fig. 6C**). Representative sections from the indicated regions are shown in **Fig. 6D**. Since TRAP recombination and c-Fos immunoreactivity may have different sensitivities to label activated neurons, we repeated this experiment in an independent cohort of TRAP2/Rosa26-LSL-tdTomato mice, but now reversed the BHB-Phe/Lac-Phe sequence. In this reversed experiment, Lac-Phe-activated neurons were identified by TRAP recombination, whereas BHB-Phe-activated neurons were identified by c-Fos immunoreactivity (**Fig. S7A**). Once again, while both Lac-Phe and BHB-Phe activated neural populations in the hypothalamus and brainstem (**Fig. S7B**), detailed analysis of each region showed that the two populations of activated neurons were largely distinct (**Fig. S7C**). We conclude that pharmacological administration of BHB-Phe activates several hypothalamic and brainstem regions implicated in feeding behaviors in a manner overlapping but distinct from that of Lac-Phe.

**Fig. 6.**
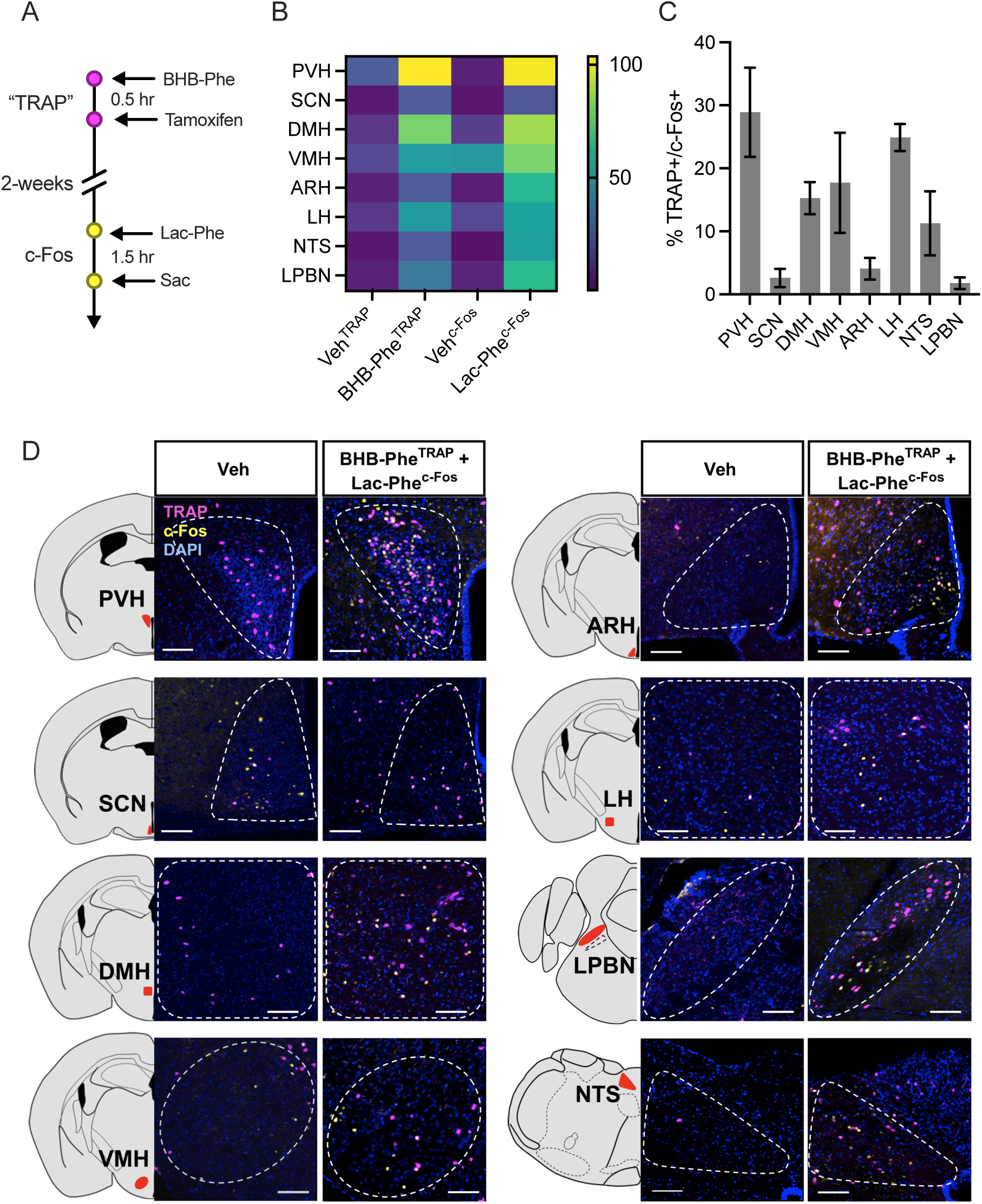
TRAP/c-Fos mapping of BHB-Phe- and Lac-Phe-activated neurons in the brain. (A) A schematic diagram of the experimental design for mapping BHB-Phe- and Lac-Phe-activated neurons. TRAP, targeted recombination in active populations. (B) Heat map showing the number of Veh^TRAP^, BHB-Phe^TRAP^, Veh^c-Fos^ and Lac-Phe^c-Fos^ labeled neurons in various brain regions. ARH, arcuate nucleus of the hypothalamus; DMH, dorsomedial hypothalamus; LH, lateral hypothalamus; LPBN, lateral parabrachial nucleus; NTS, nucleus of the solitary tract; PVH, paraventricular hypothalamus; SCN, superchiasmatic nucleus; VMH, ventromedial hypothalamus. (C,D) Quantification (C) and representative sections (D) of TRAP+/c-Fos+ neurons in the indicated brain regions. For B,C, N = 3 per group. Data are shown as mean ± SEM. Scale bars, 100 µm.

### CNDP2 and BHB-amino acids in humans

Lastly, we sought to understand the generality of the CNDP2-dependent amino acid BHB-ylation pathway to humans. First, we obtained recombinant human CNDP2, which exhibited the expected in vitro BHB-ylation activity using BHB and phenylalanine as substrates (**Fig. 7A**). Michaelis-Menten kinetics using increasing concentrations of BHB also revealed similar substrate affinity and maximal velocity to the mouse CNDP2 enzyme (**Fig. 7B**). Next, we identified three human cell lines expressing hCNDP2: U937 macrophage cells, Caco-2 gut epithelial cells, and Panc1 pancreatic ductal cells. For each human cell line, we generated control and hCNDP2-KO lines via CRISPR/Cas9. Complete loss of hCNDP2 was validated by Western blotting using our anti-CNDP2 antibody (**Fig. 7C-E**). Knockout of hCNDP2 in each cell line resulted in near complete ablation of cell lysate BHB-amino acid synthesis activity. These data show that an endogenous amino acid BHB-ylation activity is present in human cells and primarily mediated by CNDP2 (**Fig. 7C-E**). Lastly, to determine if BHB-amino acids are endogenous human metabolites, we measured plasma levels of BHB-amino acids from participants in a trial of exogenous ketone supplementation. Samples from individuals living with type 2 diabetes were from a pre-registered randomized crossover trial^29^ (NCT04194450, approved by UBC CREB H19-02947). After fasting for 24 h, participants consumed a ketone monoester drink (0.3 g/kg HVMN Ketone Ester) and plasma was collected 1 h later. BHB-Phe, BHB-Leu, BHB-Val and BHB-Met were also detectable in baseline plasma samples and elevated after ketone ester drink (**Fig. 7F**). As expected, levels of BHB were increased by the ketone ester drink while levels phenylalanine, Lac-Phe and lactate remained unchanged (**Fig. 7G**). Therefore both CNDP2-mediated amino acid BHB-ylation and BHB-amino acid metabolites are conserved in humans.

**Fig. 7.**
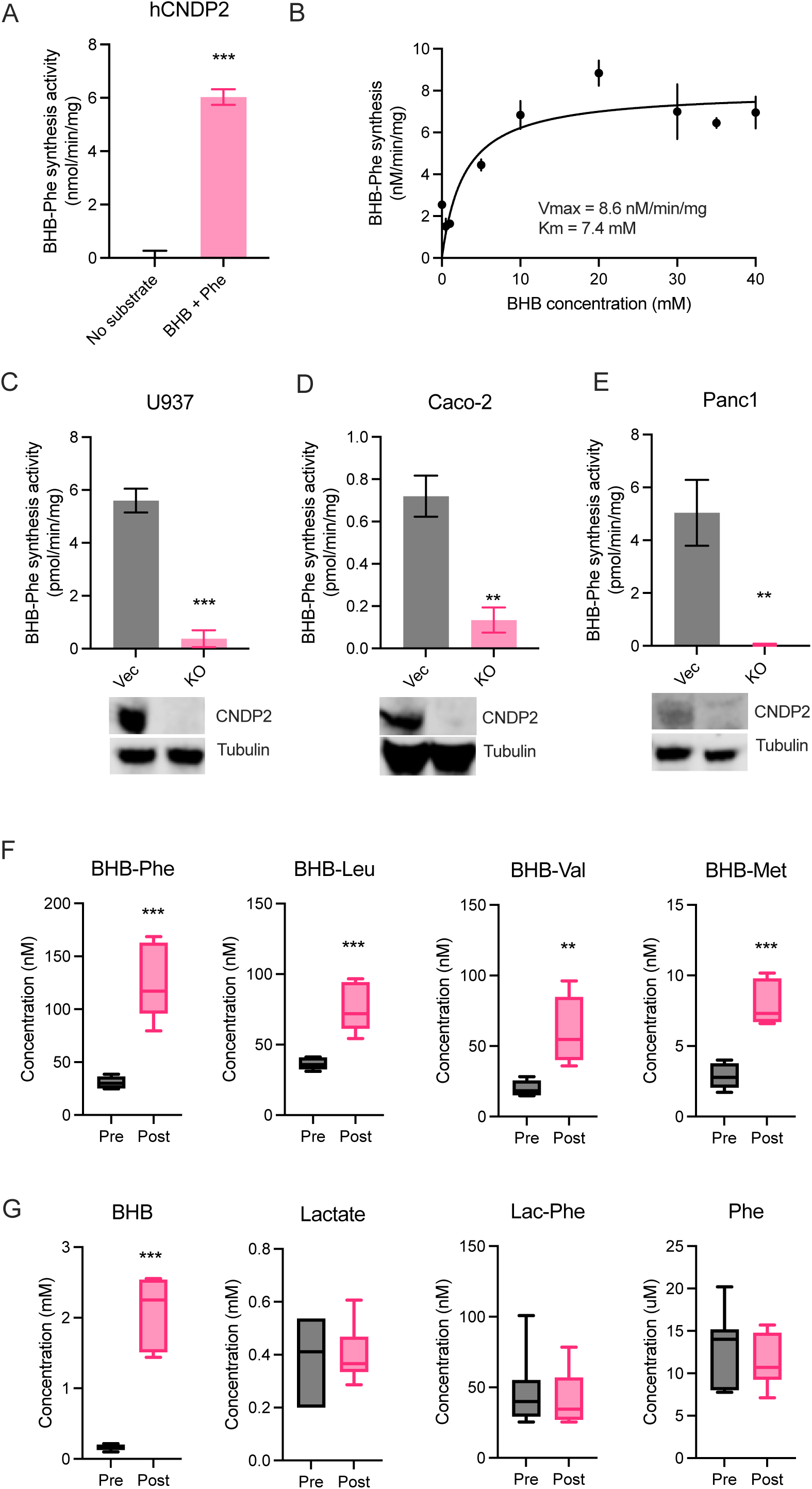
Human CNDP2 activity and BHB-amino acids in human plasma. (A,B) BHB-Phe synthetase activity of recombinant human CNDP2 provided with the indicated substrates (A) and Michaelis-Menten kinetics of recombinant human CNDP2 protein with increasing concentrations of BHB substrates (B). (C-E) Top: BHB-Phe synthesis activity of cell lysates from WT or CNDP2-KO human cell lines U937 (C), Caco-2 (D) or PANC-1 (E). Bottom: Western blot using an anti-CNDP2 (upper) or anti-tubulin (lower) antibody for WT and CNDP2-KO U937 (C), Caco-2 (D), and PANC-1 (E) cells. (F,G) Levels of BHB-amino acids (F) or the indicated metabolite (G) in human plasma at baseline or 60 minutes post ketone ester drink administration (0.3 g/kg ketone ester). For A and C-E, reactions were performed with 20 mM substrates at 37°C for 1 hour. For A-E, N=3-5/group. For F,G, N=7/group. For A-E, data are shown as mean ± SEM. For F and G, data are shown as box-and-whisker pots. P-values were calculated by Student’s two-sided t-test.

## Discussion

Here we show that CNDP2 controls a secondary pathway of BHB metabolism leading to the production of a family of BHB-derived metabolites, the BHB-amino acids. In addition, BHB-Phe, the most abundant BHB-amino acid, is a structural and functional congener of Lac-Phe. These data establish that the biochemical pathways of BHB extend beyond primary metabolic intermediates and include BHB-derived signaling metabolites that regulate energy homeostasis. That CNDP2 can accept either BHB or lactate as a substrate represents an unusual biochemical mechanism that directly couples a metabolic state with the production of bioactive metabolites. To the best of our knowledge, such a multi-functional, flux-dependent enzyme coupling mechanism has not been previously described. One interpretation of these data is that CNDP2 functions a “sensor” of glycolytic or ketosis flux, depending on whether lactate or BHB levels are elevated. Interestingly, the effector is simply a metabolic derivative of the substrate, and therefore represents a one of the simplest models by which a signal can be converted to an effector.

The chemical logic of CNDP2-dependent BHB-amino acid biosynthesis mirrors that of other signaling molecule: in every case, lower abundance bioactive species are produced from higher abundance precursors. For instance, steroid hormones are produced from cholesterol; thyroid hormones are produced from tyrosine; prostaglandins are produced from fatty acids; and histamine is produced from histidine. Our data suggests that the pool of abundant precursors is not limited to amino acids, cholesterol, or lipids, but can also include other abundant metabolic substrates such as BHB. In addition, this CNDP2-dependent mechanism is operational in *Cndp2*+ cells (e.g., macrophages, other immune cells, and epithelial cells of kidney and gut), which are cell types not classically associated with BHB metabolism.

While past studies have reported the anorexigenic and anti-obesity effects associated with elevated ketones in mice^30,31^ and in humans,^32^ this is by no means a consolidated phenomenon. Our data demonstrates that the effects of BHB also extend to BHB-derived metabolites. Therefore, potential variations in levels of BHB-amino acids may be an important contributor to the conflicting associations of ketosis and energy balance reported in previous studies. Indeed, our own data demonstrates that changes in the circulating BHB-amino acid levels are correlated to, but not linearly determined by, changes in circulating levels of BHB itself; consequently, control for variation in BHB-amino acid levels (and potentially *CNDP2* genotype) should be potentially considered in future studies of ketosis and obesity. From a teleological point of view, high ketone levels are a product of both increased hepatic ketogenesis and adipose lipolysis (to provide fatty acid substrates). Consequently, a high ketone state demonstrates that sufficient adipose lipid stores are available for ketogenesis. Therefore, one potential interpretation of the association of high ketones and food intake suppression may be that ketones (and BHB-amino acids) signal a state of fat sufficiency. In addition, our data point to hypothalamic and brainstem neural populations as potential downstream targets of BHB-Phe.

Our studies here also expand our understanding of the gene-by-environment interactions in energy balance. Previously, we had shown that CNDP2-KO mice only exhibit a body weight phenotype following treadmill running or metformin treatment, but not under “standard” high fat diet feeding conditions. Therefore phenotypes associated with the *Cndp2* gene are only revealed when the appropriate and specific environmental stimulus is provided. Our studies identify ketone ester administration and ketogenic diet as environmental contributors to CNDP2-dependent phenotype and suggest that additional nutritional or physiologic perturbations that increase ketogenesis may be relevant environmental stimuli that interact with the *Cndp2* gene.

There are two reasons why BHB-amino acids were robustly detectable in our mass spectrometry analysis, but not annotated in prior metabolomic studies. First, our enzymological studies of CNDP2, and the close chemical parallels between lactate and BHB, provided a compelling and directly testable biochemical hypothesis for the biosynthetic origins of BHB-amino acids. Second, our chemical synthesis of BHB-amino acid standards, which are otherwise not commercially available, enabled confirmation of the retention time and fragmentation of the endogenous peaks. Our strategy for detecting BHB-amino acids suggest that metabolome space might be more generally annotated by combining authentic metabolite standards with hypotheses about the chemical similarity of substrates and promiscuity of biochemical reactions.

While our studies here only examined the role of BHB-amino acids in the context of energy homeostasis, the physiologic functions of BHB-amino acids may extend to other physiologic contexts as well. For instance, ketosis is being explored in a variety of other contexts, such as in neurodegenerative diseases,^33^ inflammation,^34^ muscle resilience,^35^ cancer treatment,^36^ and several other age-associated diseases. In addition, elevated BHB is observed in other pathophysiologic conditions, such as diabetic ketoacidosis. Our data demonstrate that BHB-amino acids are also produced when levels of BHB are high, raising the possibility that the effects of ketosis and BHB in these other contexts might also be, at least in part, mediated by concomitant production of BHB-amino acids.

### Limitations of the Study

There are four main limitations of this study. First, we show that overlapping, but distinct populations of hypothalamic and brainstem neurons are activated BHB-Phe. However, their molecular identities and the functional consequences of their activation remain undetermined. Second, our gain-of-function sufficiency studies with pharmacological administration of BHB-amino acids achieved supraphysiologic levels of metabolites in circulation. While lower, more physiologically relevant concentrations were not tested here, such experiments may reveal more about the subtleties of how these ketone metabolites influence feeding and neural circuits in a physiological context. It is possible that lower, more physiologically relevant concentrations could selectively activate different subpopulations of neurons or produce more nuanced effects on feeding behavior or body weight. Such experiments might clarify the presence of potential dose-dependent “entourage effects,” where combinations of BHB-amino acids act synergistically at lower doses to modulate neural or metabolic responses. Third, we have not yet identified CNDP2 point mutants that can only accept either BHB or lactate as substrates. Such mutants which would be enable functional dissection of these multiple biochemical branches of CNDP2 activity in vitro and in vivo. Fourth, our study uses global CNDP2-KO mice which does not enable specific assignment of cell types or tissues that contribute to total BHB-amino acid synthesis in vivo.

## Acknowledgements

We thank members of the Long and Svensson laboratories for helpful discussions and Dr. Paul Emmerson (Eli Lilly & Co.) for sharing the anti-GFRAL neutralizing antibody and control IgG antibody. This work was supported by the NIH (DK124265 and DK130541 to JZL; DK125260, DK111916, and DK116074 to KJS; GM113854 to VLL; HD112123 to MW), the Phil & Penny Knight Initiative for Brain Resilience at the Wu Tsai Neurosciences Institute (research grant to J.Z.L.), the Ono Pharma Foundation (research grant to JZL), the Stanford Wu Tsai Human Performance Alliance (research grant to JZL and fellowship to XL and MDMG), the Stanford Bio-X (SIGF Graduate Student Fellowship to VLL), the Jacob Churg Foundation (research grants to JZL and KJS), the American Heart Association (fellowship #905674 to MZ), the Stanford School of Medicine (Dean’s Postdoctoral Fellowship LC), and the Fundación Alfonso Martin Escudero (fellowship to MDMG and ADG), and USDA/CRIS (51000-064-01S to YX).

## Author contributions

Conceptualization, M.D.M.G. and J.Z.L. Investigation, M.D.M.G., M.W. V.L.L., X.L., W.W., A.S.-H.T., S.H.R., L.C., M.Z., J.S., A.D.G., K.D., A.A.B., F.F.M., L.L. Writing – Original Draft, M.D.M.G., J.Z.L. Writing – Review & Editing, Y.X., J.Z.L. and M.D.M.G. Resources S.M.B., H.I., B.O., J.P.L., C.W., C.D.G., A.L., E.L.G., Y.D. Supervision J.Z.L., Y.X. S.M.B., and K.J.S.

## Quantification and statistical analysis

Statistical analysis was performed in Prism 10.2.3. All data was expressed as mean ± SEM unless otherwise specified. A student’s two-sided t-test was used for pair-wise comparisons. Two-way ANOVA with repeated measures in one factor were used for time course data of repeated measurements. Unless otherwise specified, statistical significance was set as P < 0.05. The specific test, P value symbol and error bar meaning, definition of center, and number of replicates are noted in figure legends.

## Supplementary Information

**Data S1.** ^1^H NMR spectra for all BHB-amino acids, related to STAR Methods.

**Table S1.** Multiple reaction monitoring transitions for measuring endogenous metabolites.

## References

1. Newman, J. C. & Verdin, E. β-Hydroxybutyrate: A Signaling Metabolite. Annu. Rev. Nutr. 37, 51–76 (2017).

2. Puchalska, P. & Crawford, P. A. Metabolic and Signaling Roles of Ketone Bodies in Health and Disease. Annu. Rev. Nutr. 41, 49–77 (2021).

3. Rahman, M. et al. The b-hydroxybutyrate receptor HCA 2 activates a neuroprotective subset of macrophages. Nat. Commun. 5, 1–11 (2014).

4. Kimura, I. et al. Short-chain fatty acids and ketones directly regulate sympathetic nervous system via G protein-coupled receptor 41 (GPR41). Proc. Natl. Acad. Sci. U. S. A. 108, 8030–8035 (2011).

5. Xie, Z. et al. Metabolic Regulation of Gene Expression by Histone Lysine β-Hydroxybutyrylation. Mol. Cell 62, 194–206 (2016).

6. Shirakawa, K. et al. Suppression of Oxidative Stress by beta-Hydroxybutyrate, an Endogenous Histone Deacetylase Inhibitor. Science (80-.). 339(6116), 211–214 (2013).

7. Bueno, N. B., De Melo, I. S. V., De Oliveira, S. L. & Da Rocha Ataide, T. Very-low-carbohydrate ketogenic diet v. low-fat diet for long-term weight loss: A meta-analysis of Randomised controlled trials. Br. J. Nutr. 110, 1178–1187 (2013).

8. Soto-Mota, A., Norwitz, N. G., Evans, R., Clarke, K. & Barber, T. M. Exogenous ketosis in patients with type 2 diabetes: Safety, tolerability and effect on glycaemic control. *Endocrinol*. Diabetes Metab. 4, 1–7 (2021).

9. Weber, D. D. et al. Ketogenic diet in the treatment of cancer – Where do we stand? Mol. Metab. 33, 102–121 (2020).

10. Goldberg, E. L., Letian, A., Dlugos, T., Leveau, C. & Dixit, V. D. Innate immune cell-intrinsic ketogenesis is dispensable for organismal metabolism and age-related inflammation. J. Biol. Chem. In press (2023) doi:10.1016/j.jbc.2023.103005.

11. Pierre, K. et al. Enhanced expression of three monocarboxylate transporter isoforms in the brain of obese mice. J. Physiol. 583, 469–486 (2007).

12. Shafqat, N. et al. A structural mapping of mutations causing succinyl-CoA:3-ketoacid CoA transferase (SCOT) deficiency. J. Inherit. Metab. Dis. 36, 983–987 (2013).

13. Wang, W. et al. A PRDM16-Driven Metabolic Signal from Adipocytes Regulates Precursor Cell Fate. Cell Metab. (2019) doi:10.1016/j.cmet.2019.05.005.

14. Cheng, C. W. et al. Ketone Body Signaling Mediates Intestinal Stem Cell Homeostasis and Adaptation to Diet. Cell 178, 1115–1131.e15 (2019).

15. Karagiannis, F. et al. Impaired ketogenesis ties metabolism to T cell dysfunction in COVID-19. Nature 609, 801–807 (2022).

16. Jansen, R. S. et al. N -lactoyl-amino acids are ubiquitous metabolites that originate from CNDP2-mediated reverse proteolysis of lactate and amino acids . Proc. Natl. Acad. Sci. (2015) doi:10.1073/pnas.1424638112.

17. Li, V. L. et al. An exercise-inducible metabolite that suppresses feeding and obesity. Nature 606, 785–790 (2022).

18. Xiao, S. et al. Lac-Phe mediates the effects of metformin on food intake and body weight. Nat. Metab. 2024 1–11 (2024) doi:10.1038/s42255-024-00999-9.

19. Scott, B. et al. Metformin and feeding increase levels of the appetite-suppressing metabolite Lac-Phe in humans. Nat. Metab. 15497, (2024).

20. Schaum, N. et al. Single-cell transcriptomics of 20 mouse organs creates a Tabula Muris. Nature 562, 367–372 (2018).

21. Smith, C. A. et al. METLIN: a metabolite mass spectral database. Ther. Drug Monit. 27, 747–751 (2005).

22. Wishart, D. S. et al. HMDB 5.0: The Human Metabolome Database for 2022. Nucleic Acids Res. 50, D622–D631 (2022).

23. Wang, M. et al. Sharing and community curation of mass spectrometry data with Global Natural Products Social Molecular Networking. Nat. Biotechnol. 34, 828–837 (2016).

24. Gall, W. E. et al. Α-Hydroxybutyrate Is an Early Biomarker of Insulin Resistance and Glucose Intolerance in a Nondiabetic Population. PLoS One 5, (2010).

25. Nilsen, M. S. et al. 3-Hydroxyisobutyrate, A Strong Marker of Insulin Resistance in Type 2 Diabetes and Obesity That Modulates White and Brown Adipocyte Metabolism. Diabetes 69, 1903–1916 (2020).

26. Emmerson, P. J. et al. The metabolic effects of GDF15 are mediated by the orphan receptor GFRAL. Nat. Med. 23, 1215–1219 (2017).

27. Worth, A. A. et al. The cytokine GDF15 signals through a population of brainstem cholecystokinin neurons to mediate anorectic signalling. Elife 9, 1–19 (2020).

28. DeNardo, L. A. et al. Temporal evolution of cortical ensembles promoting remote memory retrieval. Nat. Neurosci. 22, 460–469 (2019).

29. Monteyne, A. J. et al. A ketone monoester drink reduces postprandial blood glucose concentrations in adults with type 2 diabetes: a randomised controlled trial. Diabetologia 67, 1107–1113 (2024).

30. Nasser, S. et al. Ketogenic diet administration to mice after a high-fat-diet regimen promotes weight loss, glycemic normalization and induces adaptations of ketogenic pathways in liver and kidney. Mol. Metab. 65, 101578 (2022).

31. Laeger, T., Metges, C. C. & Kuhla, B. Role of β-hydroxybutyric acid in the central regulation of energy balance. Appetite 54, 450–455 (2010).

32. Stubbs, B. J. et al. A Ketone Ester Drink Lowers Human Ghrelin and Appetite. Obesity 26, 269–273 (2018).

33. Fortier, M. et al. A ketogenic drink improves cognition in mild cognitive impairment: Results of a 6-month RCT. Alzheimer’s Dement. 17, 543–552 (2021).

34. Youm, Y. H. et al. The ketone metabolite β-hydroxybutyrate blocks NLRP3 inflammasome-mediated inflammatory disease. Nat. Med. 21, 263–269 (2015).

35. Benjamin, D. I. et al. Fasting induces a highly resilient deep quiescent state in muscle stem cells via ketone body signaling. Cell Metab. 34, 902–918.e6 (2022).

36. Dmitrieva-Posocco, O. et al. β-Hydroxybutyrate suppresses colorectal cancer. Nature 605, 160–165 (2022).

37. Unno, H. et al. Structural basis for substrate recognition and hydrolysis by mouse carnosinase CN2. J. Biol. Chem. 283, 27289–27299 (2008).

38. Allouche, A. Software News and Updates Gabedit — A Graphical User Interface for Computational Chemistry Softwares. J. Comput. Chem. 32, 174–182 (2012).

39. Pettersen, E. F. et al. UCSF Chimera - A visualization system for exploratory research and analysis. J. Comput. Chem. 25, 1605–1612 (2004).

